# Complementary and divergent roles for Ctage5 and Tango1 in zebrafish

**DOI:** 10.1101/2020.04.30.070664

**Authors:** Eric M. Clark, Brian A. Link

**Affiliations:** Department of Cell Biology, Neurobiology and Anatomy, Medical College of Wisconsin, Milwaukee, WI, United States of America

**Keywords:** COPII, Secretion, Collagen, Lipoprotein, ER Stress, Zebrafish

## Abstract

Coat protein complex II (COPII) factors mediate cargo export from the endoplasmic reticulum (ER), but bulky collagens and lipoproteins are too large for traditional COPII vesicles. Mammalian CTAGE5 and TANGO1 have been well characterized individually as specialized cargo receptors at the ER that function with COPII coats to facilitate trafficking of bulky cargoes. Here, we present a genetic interaction study in zebrafish of deletions in *ctage5*, *tango1*, or both to investigate their potential distinct and complimentary functions. We found that Ctage5 and Tango1 have different roles related to organogenesis, collagen versus lipoprotein trafficking, stress-pathway activation, and survival. While deletion of both *ctage5* and *tango1* compounded phenotype severity, deletion of either factor alone revealed novel tissue specific defects in the building of heart, muscle, lens, and intestine, in addition to the previously described roles in the development of neural and cartilage tissues. Together, our results suggest that Ctage5 and Tango1 have overlapping, but also divergent roles in tissue development and homeostasis.

**Summary:** In this genetic study Ctage5 and Tango1 endoplasmic reticulum cargo receptors were investigated together *in vivo* for the first time. Cell differentiation, survival, trafficking, and stress pathway activation were investigated.

## Introduction

Secretory proteins are trafficked by coat protein complex II (COPII) vesicles that form at specialized sites of the endoplasmic reticulum (ER) called ER-exit sites (ERES) (Bonifacino and Glick, 2004; Lee et al., 2004). These sites contain conserved proteins like SEC12, SEC16, SEC13, SEC31, SEC23, SEC24, SAR1 among others that are important for COPII-mediated trafficking (Saito et al., 214.; Watson et al., 2006; Maeda et al., 2017; Townley et al., 2008; Kim et al., 2012; Sato & Nakano, 2005). Some bulky molecules are too large to fit into traditional COPII vesicles which are 60-90nm (Malhotra and Erlman, 2015; Miller and Schekman, 2013). Recently, a transport system using specialized cargo receptors, the TANGO1(MIA3)/CTAGE5(MEA6) family, has been described for large molecules like collagens, chylomicrons, and very low density lipoproteins (VLDLs) (Saito et al., 2011; Santos et al., 2016; Wilson et al, 2011; Saito et al., 2009; Maiers et al., 2016; Rios-Barrera et al., 2017; Maeda et al., 2016). Members of the TANGO1/CTAGE5 family form a ring-like structure at the base of ERES vesicles and promote their expansion to accommodate large cargo (Raote et al., 2017; Lui et al., 2017; Reynolds et al. 2019) or potentially facilitate tubular connections for transfer of material to the cis-Golgi (Kurokawa et al., 2014; Raote and Malhotra, 2019).

The TANGO1/CTAGE5 family is defined by shared protein domains. CTAGE5 contains a transmembrane motif and cytoplasmic domains including two coiled coil regions that bind to TANGO1 and SEC12, and a proline rich domain that binds to COPII inner coat membrane proteins SEC23 and SEC24 (Saito et al., 2011; Saito et al., 2014). Conversely, MIA2 contains only a luminal SH3 domain used for binding cargo (Pitman et al., 2011). TANGO1-LIKE (TALI) is a chimeric protein produced by a fusion transcript of MIA2 and CTAGE5 (Pitman, 2011). Through alternate transcriptional initiation, the *TANGO1* locus produces two protein isoforms (Maeda et al., 2016). TANGO1L has similar domains to TALI including a luminal cargo binding domain, coiled coil domains, and a proline rich domain (Bard et al., 2006; Saito et al., 2009). TANGO1S is a short isoform that contains only the cytoplasmic domains and is analogous to cTAGE5 (Maeda et al., 2016).

In vitro studies have characterized TANGO1/CTAGE5 family member functional domains for ERES complex formation, cargo binding, and trafficking (Saito et al., 2009; Saito et al., 2011; Ma & Goldberg, 2016; Maeda et al., 2016; Santos et al., 2016; Maiers et al., 2016; Yuan et al., 2018; Maeda et al., 2019). Collagen is the most abundant protein in the body and crucial for the extracellular matrix. For collagen VII trafficking, TANGO1S and TANGO1L are individually important for export from the ER in HSC-1 cells (Saito et al., 2009; Maeda et al., 2016), but are exchangeable as overexpression of either isoform was able to rescue collagen VII defects from loss of the other (Maeda et al., 2016). CTAGE5 is also necessary for collagen VII secretion (Saito et al., 2011) suggesting that these factors have overlapping functions. However, loss of TANGO1, but not TALI blocked collagen XII export in Caco2 cells indicating there are potential differences for how CTAGE5/TANGO1 family members export specific collagens in specific cell types (Santos et al., 2016).

In vivo studies of TANGO1/CTAGE5 family members have been relatively limited, likely because in mice *Tango1* deletion is perinatally lethal (Wilson et al., 2011) and *Ctage5* deletion is essential for early embryonic development (Wang et al., 2016). *Tango1l* deletion caused buildup of multiple collagen sub-types in chondrocytes, fibroblasts, endothelial cells, and mural cells (Wilson et al., 2011). Conditional knockout of *Ctage5* in neurons of mice show that in addition to secretory components, non-secretory protein trafficking is also affected (Zhang et al., 2018). Similarly, *Drosophila* studies have showed secretion of multiple extracellular proteins including mucins, laminins, and perlecan are disrupted after loss of Tango1 (Petley-Ragan et al., 2016; Liu et al., 2017; Rios-Barrera et al. 2017; Reynolds et al., 2019). Of significance, however, studies of *Ctage5* and *Tango*1 deletions within a single animal model have not been reported, highlighting a need to investigate their potential redundancies or unique functions in different cell types.

Zebrafish have previously been used to study COPII proteins, including the *feelgood* (*creb3l2*) (Melville et al., 2011), and *crusher* (*sec23*) (Lang et al., 2006) mutants, but neither Ctage5 nor Tango1 have been investigated in this model system. In this study, we generated *ctage5* and *tango1* mutant zebrafish and characterized phenotypes in the single as well as double homozygous mutant animals. We find that like other COPII pathway mutants, *ctage5* and *tango1* mutant zebrafish are small and have decreased survival. Overall, our studies reveal that Ctage5 and Tango1 have complimentary, but also divergent roles in craniofacial development, collagen and lipoprotein trafficking, and in activation of cellular stress pathways.

## Results

### Morphogenesis and survival are differentially affected by *ctage5* or *tango1l* deletion

To investigate how loss of *ctage5* and *tango1* deletion affects zebrafish development and survival, large portions of genomic coding sequence were deleted with Crispr/Cas9, eliminating critical functional domains (Fig. 1A). Based on Ensembl transcript analysis, the zebrafish *mia2* gene locus only transcribes *ctage5*. Transcripts of the deletion variant are decreased two-fold compared to wild-type *ctage5* mRNA. For the *mia3* gene locus, the deletion completely eliminates *tango1l*, but spares *tango1s*. Primers designed to measure *tango1l* and *tango1s* transcript levels did not reveal a change between wild-type and *tango1* mutant embryos, suggesting that Tango1s protein could still be made. Our analysis indicated there was no compensatory upregulation of *ctage5* or *tango1* expression with deletion of the other gene (Fig. S1).

**Fig 1:**
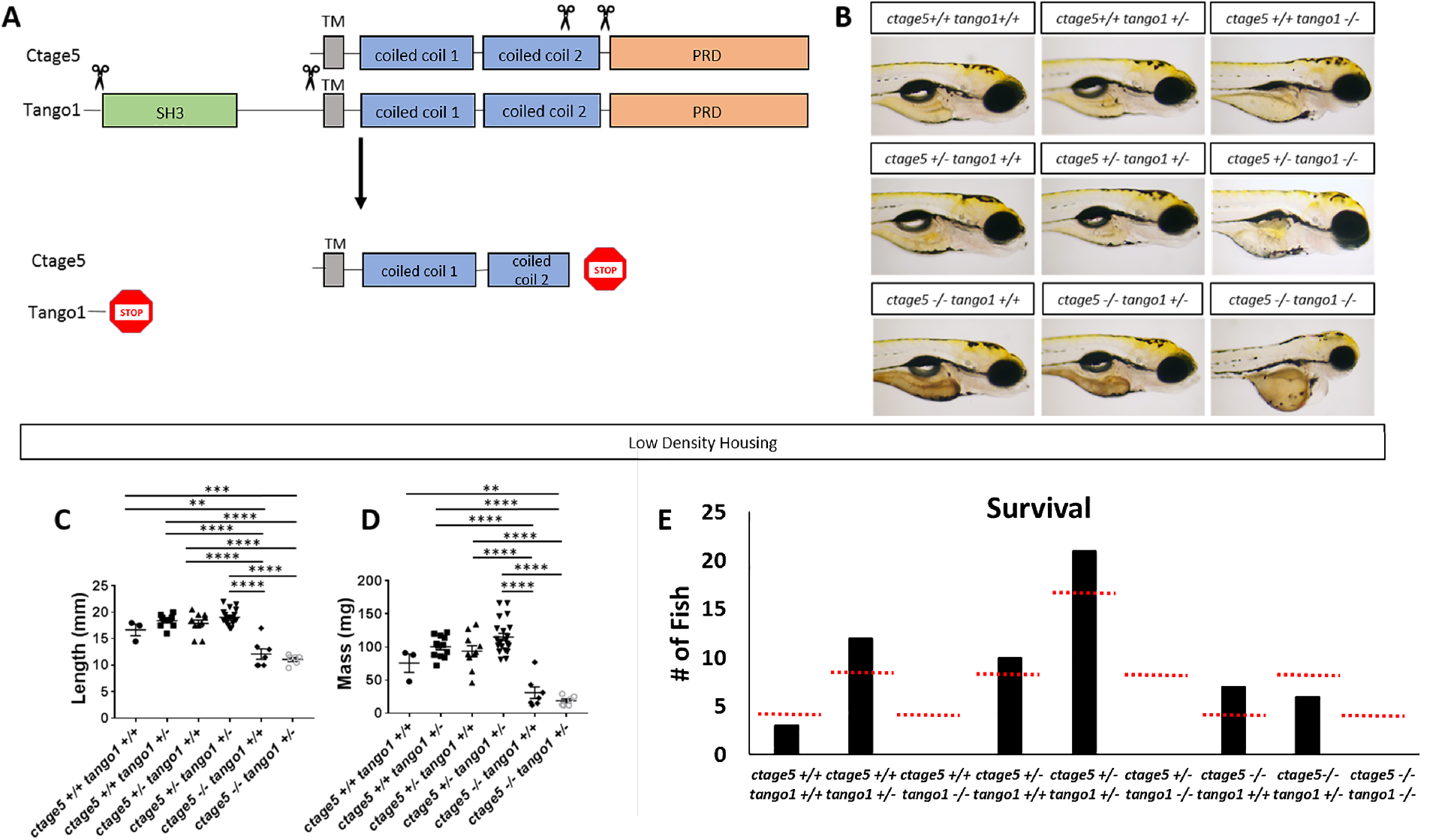
*ctage5* and *tango1* mutations affect size and survival of zebrafish: (A) Schematic showing CRISPR guide RNA cut sites (scissors) and the estimated final truncated proteins. (B) Representative brightfield images of all combinations of *ctage5* and *tango1* mutations in 4 dpf zebrafish larvae. (C,D,E) Length (C; one-way ANOVA, F=38.6, p<0.0001), mass (D; one-way ANOVA, F=28.22, p<0.0001) and survival (E; χ^2^=24.59, p<0.05, n=59), measurements of all combinations of *ctage5* and *tango1* mutant zebrafish at 2 months raised in a low density environment (about 10 fish per tank). Red-dotted lines in (E) represent expected survival, and black bars are actual survival. **=p<0.01, ***=p<0.001, ****= p<0.0001.

Overtly, *ctage5* mutants look normal except for having darker yolk (Fig. 1B). However, *tango1* mutant larvae are significantly shorter and have obvious craniofacial defects as the mouth fails to protrude past the eyes (Fig. 1B, S2A). *ctage5;tango1* double mutants have similar, but more severe phenotypes compared to single mutants, including some unique characteristics such as smaller eyes (Fig. 1B). Although body length of larval *ctage5* mutants is normal, those that survived due to low density rearing (10 fish per tank) show reduced size as adults (Fig. 1C-E). When raised at high-density (30 fish per tank), mutants had reduced survival (Fig. S2B). Regardless of rearing density, most *ctage5* mutants die by 1 year, while heterozygous and wild-type siblings show normal survival past 3 years. Together, these results suggest that *ctage5* and *tango1* deletions differentially affect zebrafish during development, pointing to potential differences in cellular function.

### Craniofacial defects in *tango1* mutants are accentuated with loss of *ctage5*

Mutations in the COPII trafficking pathway commonly cause craniofacial defects (Lang et al., 2006; Townley et al., 2008; Melville et al., 2011). For example, in mice loss of Tango1 causes dwarfed embryos with defects in cartilage formation and bone mineralization (Wilson et al., 2011). Alcian blue staining confirmed that like mice, zebrafish *tango1* mutant embryos also have defects in cartilage formation at 4 dpf (Fig. 2A). *ctage5;tango1* double mutants have a near complete absence of cartilage staining (Fig. 2A). Histology also revealed severe craniofacial defects in *tango1* mutant embryos that were exacerbated by *ctage5* deletion (Fig. 2B). Interestingly, ocular lens defects exemplified by increased cell death and a degenerative lens were only observed in *ctage5;tango1* double mutant embryos (Fig. 2C,D). Although *ctage5* mutant lenses appear normal during development, by two months lens epithelial defects were observed (Fig. 2E), suggesting a lens homeostasis role for Ctage5. Cardiac chamber dysgenesis was another unique phenotype to *ctage5;tango1* double mutants. While both *ctage5* and *tango1* mutants showed normal heart development, severe cardiac structural changes were apparent by 4dpf when these genes were disrupted in tandem (Fig. 2F). However, blood clotting in the caudal vein was noticed in both *tango1* single and *ctage5;tango1* double mutant larvae, suggesting that *tango1* deletion could be driving cardiovascular defects which only becomes obvious with the additional deletion of *ctage5* (Fig. 2G). These results extend previous *in vivo* observations demonstrating Tango1 plays a critical role in maintaining cartilage formation and overall craniofacial development (Wilson et al., 2011). However, Ctage5 also acts in this process as craniofacial defects observed with deletion of *tango1* are enhanced by *ctage5* deletion.

**Fig. 2:**
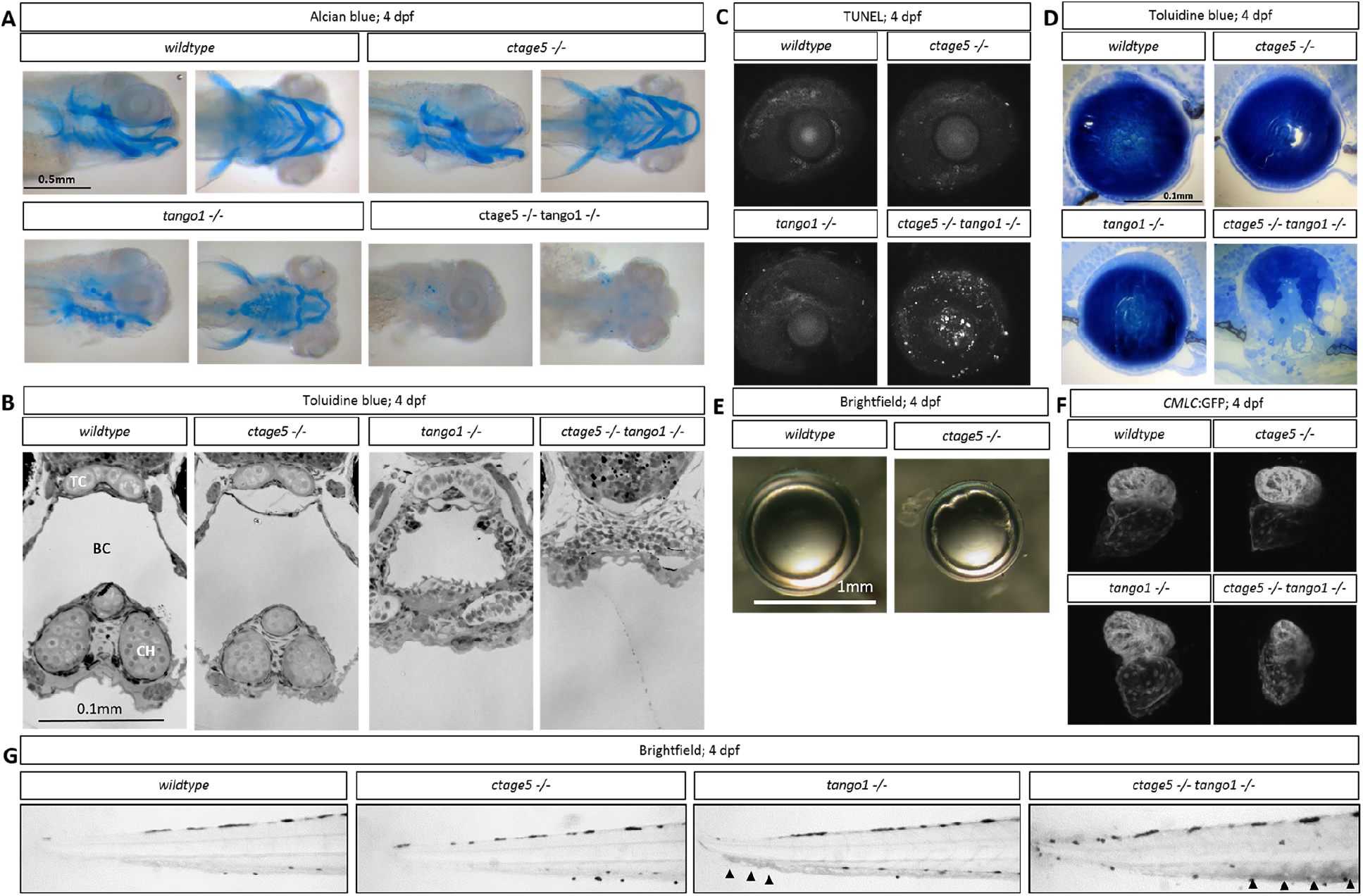
Craniofacial defects in *tango1* mutants are accentuated with *ctage5* deletion. (A) Representative lateral (left) and ventral (right) images of alcian blue stained 4 dpf larvae. (B) Representative images of plastic sectioned and toluidine blue stained 4 dpf larvae cropped to show craniofacial alterations. TC=trabeculae, BC=buccal cavity, CH=ceratohyal cartilage (C) Representative maximum intensity projection images showing TUNEL staining of 4 dpf lens and sclera. (D) Representative images of plastic sectioned and toluidine blue stained 4 dpf lens. (E) Representative images of lens dissected from 3-month-old zebrafish. (F) Representative images showing ventricle-enriched cardiac myosin light chain (*CMLC*:GFP) expression. (G) Representative brightfield image of the trunk for each genotype. Arrowheads detail blood pooling in the caudal vein.

### Large molecule trafficking differs between *ctage5* and *tango1* mutants

TANGO1 has been well established as a cargo receptor for collagen trafficking in various cell types in vitro and in vivo (Saito et al., 2009; Wilson et al., 2011; Maeda et al., 2016; Maiers et al., 2016; Rios-Barrera et al., 2017; Gorur et al., 2017). CTAGE5 is necessary for collagen VII trafficking in an epidermal carcinoma human cell line (Saito et al., 2011), but its role in secreting other collagens and in other cell lines or in vivo is still largely unknown. In vitro, lipid trafficking was also found affected after loss of TALI and TANGO1 (Santos et al., 2016). In vivo, conditional knockout of *Ctage5* in the mouse liver and nervous system significantly affected the lipidomic profile within serum and brain extracts, respectively (Wang et al., 2016; Zhang et al., 2018). To investigate large molecule trafficking in zebrafish, we performed wholemount staining for Collagen II. The branchial arch cartilage of wild-type animals showed extracellular Collagen II surrounding chondrocytes. In contrast, in both *ctage5* and *tango1* mutants, Collagen II was found in intracellular puncta within chondrocytes (Fig. 3A). With decreased trafficking of large molecules, we hypothesized that protein buildup occurred within the ER, which would be consistent with previous reports of distended ER after loss of Tango1 (Wilson et al., 2011; Petley-Ragan et al., 2016; Rios-Barrerra et al., 2017; Maiers et al., 2016). Indeed, we observed distended ER in *tango1* and double mutant chondrocytes, while the ER in *ctage5* mutants and wild-type chondrocytes was normal (Fig. 3B). We next investigated lipid trafficking using a lipid analog, bodipy-c12 that has previously been shown to require β-lipoproteins for trafficking from the yolk into the zebrafish vasculature (Miyares et al., 2014). Efficient β-lipoprotein trafficking was observed in wild-type larvae by monitoring bodipy-c12 vascular fluorescence following injection into the yolk. However, bodipy-c12 trafficking was decreased after *tango1* deletion, and nearly undetectable in *ctage5* or double mutant embryos (Fig. 3C). Oil Red O staining confirmed this significant loss of serum lipids in *ctage5* single and *ctage5*;*tango1* double mutant embryos (Fig. 3D). Together these results suggest that lipoprotein trafficking is significantly affected after deletion of Ctage5, and to a lesser extent with loss of Tango1. Conversely, in cell types that secrete large amounts of collagen, like chondrocytes, deletion of *tango1*, but not *ctage5* severely affected protein export and ER maintenance.

**Fig. 3.**
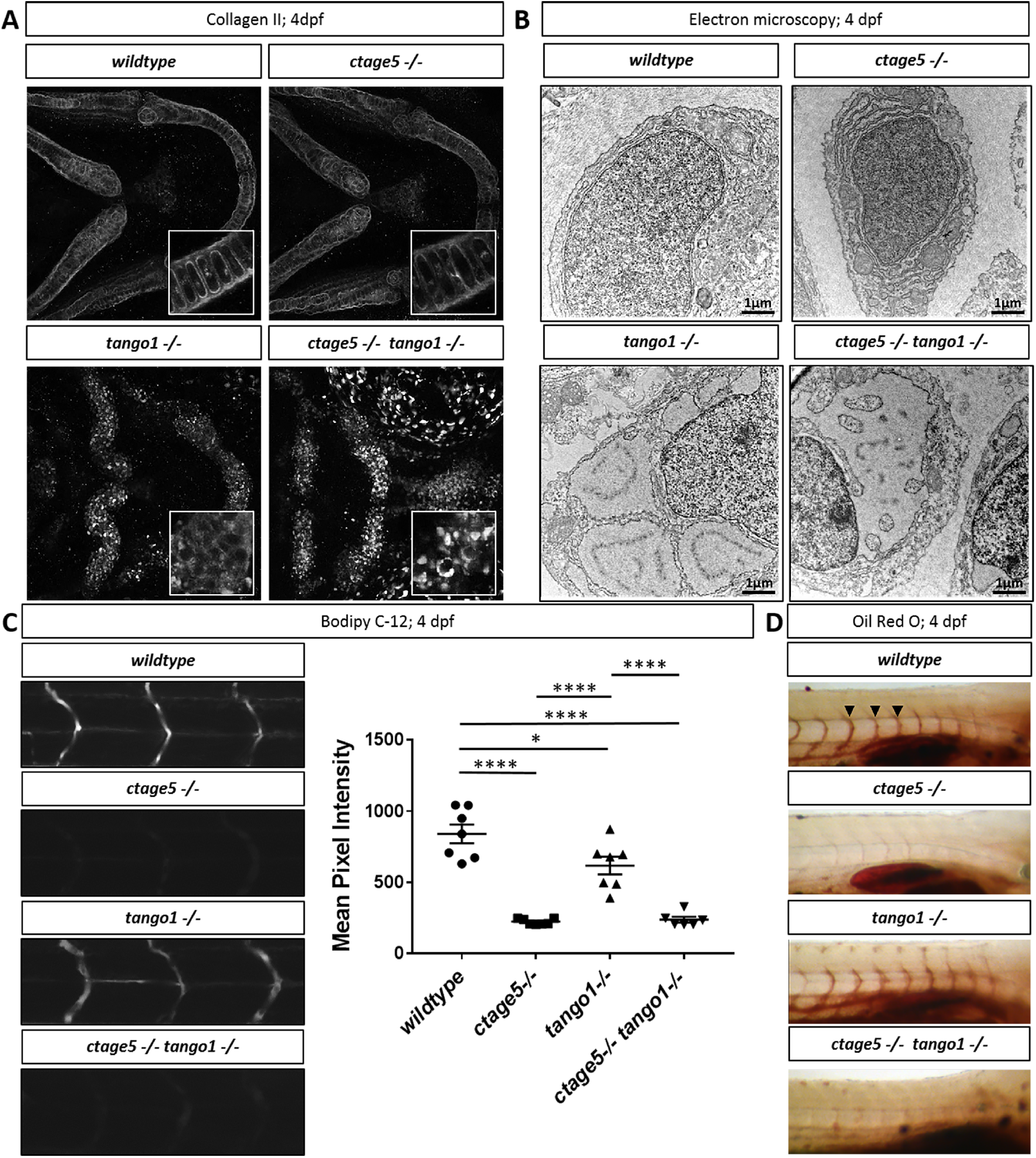
Trafficking differences in *ctage5* vs. *tango1* mutant embryos. (A) Representative 40X images showing collagen 2 wholemount immunohistochemistry in 4 dpf larvae chondrocytes. Inserts are 40X images with 3X zoom and cropped to show higher resolution chondrocyte collagen 2 staining. (B) Representative electron microscopy images of 4dpf larvae chondrocytes. (C) Representative images and quantification of Bodipy C-12 fluorescence in the intersegmental vessels (arrowheads) (One-way ANOVA, F=39.36, p<0.0001). (D) Representative images of oil red o staining for lipids in 4dpf zebrafish larvae. *=p<.05, ****= p<0.0001.

### Cell stress is differentially activated in tissues after *ctage5* or *tango1* deletion

When cellular trafficking is affected and cargo builds up at the ER, the evolutionarily conserved unfolded protein response (UPR) can be activated (Walter and Ron, 2011). This coordinated response has three major branches, the ATF6, PERK, and XBP1 pathways (Walter and Ron, 2011). After TANGO1 knockdown, XBP1 and ATF4 ER stress pathways were activated, but ATF6 was not investigated (Maiers et al., 2016; Petley-Ragan et al., 2016). After loss of TALI or TANGO1 in HEPG2 or Caco-2 cells, APOB accumulates in cells and is targeted for lysosomal degradation (Santos et al., 2016). Of further relevance for ER stress response, CTAGE5 was shown as essential for starvation-induced autophagosome formation in HeLa cells, acting at the ERES for membrane donation (Ge et al., 2017). The role of TANGO1, however, was not investigated in that study. Collectively, these reports suggest ER stress and autophagy could be altered after *CTAGE5* or *TANGO1* knockout. We recently described a transgenic zebrafish line to measure the ATF6 branch of the ER stress network (Clark et al., 2020). Using these transgenic fish, we observed that *ctage5*−/−; *tango1*−/− larvae have significantly increased *ATF6* reporter fluorescence in the spinal cord at 4 dpf (Fig. 4A, A’). As assessed by nuclei packing and cell body morphology, spinal cord cells in the double mutants are disorganized and in some instances mislocalized from the normal lamination pattern (Fig. 4B, B’, asterisk). Although spinal cord cells of *ctage5* single mutants were unaffected at 4 dpf, by 7 dpf these larvae had significantly increased *ATF6* reporter activation (Fig. 4C, C’). Furthermore, 7 dpf *ctage5* mutant larvae had increased lysosomal volume in the spinal cord and brain compared to wild-type or *tango1* mutants (Fig. 4D, D’, E, E’). In addition, within the brain, *ctage5*;*tango1* double mutant embryos showed significant increase in cell death markers, while single mutations were similar to wild-type brains (Fig. 4F, F’). As adults, *ctage5* mutant zebrafish show elevated *gap43*:GFP expression (Fig. 4G). Expression of *gap43* is known to be induced with neuronal stress (Skene, 1989; Benowitz and Routtenberg, 1997; Bormann et al., 1998; Kaneda et al., 2008; Diekmann et al., 2015). Therefore, *ctage5* mutant larvae undergo cellular stress throughout the nervous system, as evidenced by ER stress, altered cell morphology, elevated autophagy and apoptosis, which continues into adulthood. Our observations are consistent with the axonal trafficking defects described in mice with conditional *Ctage*5 deletion from the nervous system (Zhang et al., 2018). We also noticed ER stress in other tissues including the jaw in *tango1* mutant and double mutant larvae, but not in *ctage5* mutant embryos, consistent with craniofacial defects observed in Fig. 2 (Fig. 5A, A’, B, B’). Conversely, ER stress was observed in the intestine of *ctage5* mutant larvae but not *tango1* mutants at 7dpf (Fig. 5C, C’). Within trunk skeletal muscle, both *ctage5* and *tango1* mutant embryos showed ER stress (Fig. 5D, D’, E, E’). Interestingly, however, *ctage5* mutants show significantly elevated *ATF6* activation in the muscle fibers, whereas *tango1* ER stress is activated along cells lining the chevron shaped muscle segments. The role for Ctage5 or Tango1 in muscle tissue has not been previously investigated, but these data suggest Ctage5/Tango1 family of proteins also have divergent functions in that tissue.

**Fig. 4:**
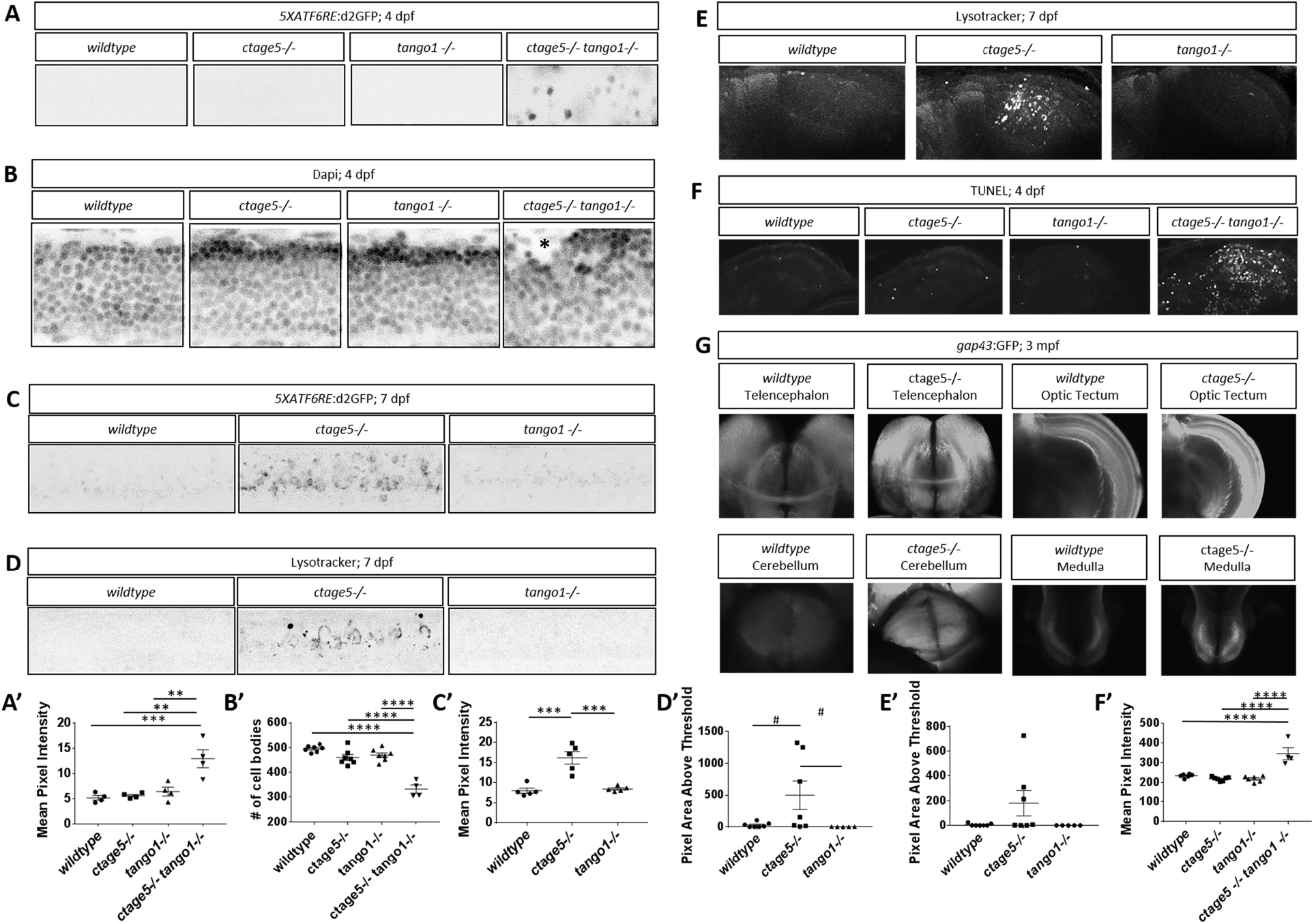
Stress pathway activation in the nervous system is specific for *ctage5* −/− and amplified by *tango* mutation. (A, A’) Representative images (A) and quantification (A’) of *5XATF6RE*:d2GFP activation in 4 dpf spinal cords (one-way ANOVA, F= 12.42, p=0.0005). (B, B’) Representative images of Dapi staining (B) and quantification of brightfield images (B’) in 4 dpf spinal cords to show cellular organization (asterisk shows cells mislocalized from the normal lamination pattern) (one-way ANOVA, F=37.51, p<0.0001). (C, C’) Representative images (C) and quantification (C’) of *5XATF6RE*:d2GFP activation in 7 dpf spinal cords (one-way ANOVA, F=22.22, p<0.0001). (D, D’) Representative images (D) and quantification (D’) of the lysotracker staining area above threshold for acidic compartments in 7 dpf spinal cords (one-way ANOVA, F=3.881, p=0.0423). (E, E’) Representative images (E) and quantification (E’) of the lysotracker staining area above threshold of lysotracker for acidic compartments in 7 dpf brains (one-way ANOVA, F=2.479, p=.1154). (F, F’) Representative images (F) and quantification (F’) of TUNEL staining for cell death in 4 dpf brains (one-way ANOVA, F=24.33, p<0.0001). (G) Representative images of *gap43*:GFP in 3-month old PACT cleared *ctage5* wildtype (WT) or homozygous mutant brains. #=p<.08, **=p<0.01, ***=p<0.001, ****= p<0.0001.

**Fig. 5:**
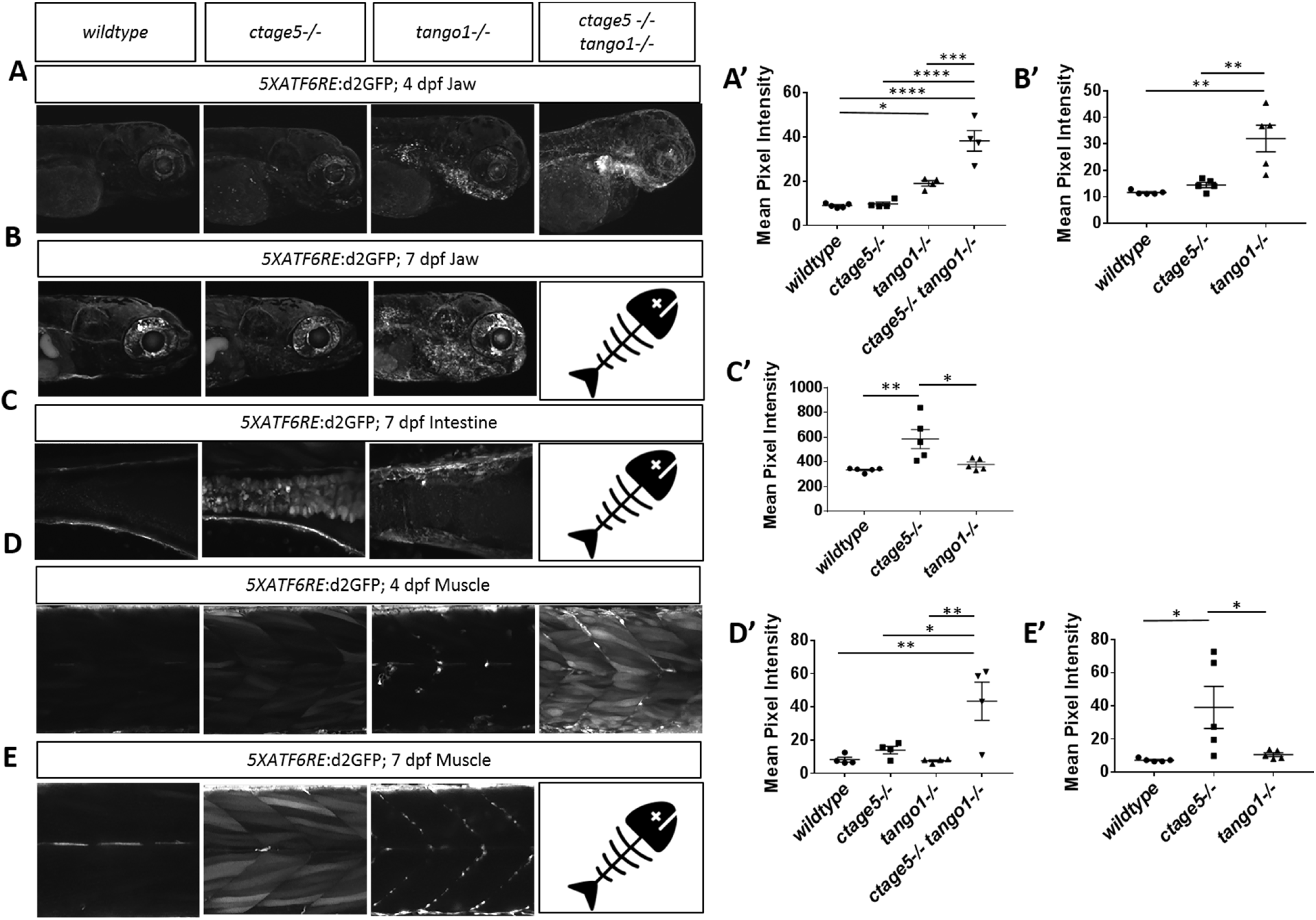
ER stress is prevalent in tissues outside of the nervous system. Representative images and quantification of 5XATF6RE:d2GFP ER stress in the jaw at 4dpf (A, A’; one-way ANOVA, F=34.44, p<0.0001) and 7dpf (B, B’; one-way ANOVA, F=13.86, p<0.0008), intestine at 7dpf (C, C’; one-way ANOVA, F=8.358, p=0.0053) and muscle at 4 dpf (D, D’; one-way ANOVA, F=8.205, p=0.0031) and 7 dpf (E, E’: F=5.624, p=0.0189) and. *=p<.05, **=p<0.01, ***=p<0.001, ****= p<0.0001. 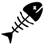 = indicates *ctage5−/− tango1−/−* mutants do not survive to 7dpf.

## Discussion

CTAGE5 and TANGO1 are both known to act in the COPII trafficking pathway (Saito et al., 2009; Saito et al., 2011), but until now the effects of deleting these proteins in the same model system has not been investigated. Our results suggest that zebrafish Ctage5 and Tango1 have complimentary, but also divergent roles across multiple tissues (Fig. 6). For example, we found that both factors affect fish growth when deleted. *tango1* mutant fish are shorter at larvae stages, whereas *ctage5* mutant larvae are morphologically normal initially, but those that survive are much smaller than their wild-type siblings. Consistent with our results, global knockout of *Tango1* inhibits growth in mouse embryos (Wilson et al., 2011). Interestingly, conditional knockout of *Ctage5* in the mouse brain or liver also results in decreased animal size (Wang et al., 2016; Zhang et al., 2018), indicating Ctage5 function in multiple organ systems is important for embryonic growth and development.

**Fig. 6:**
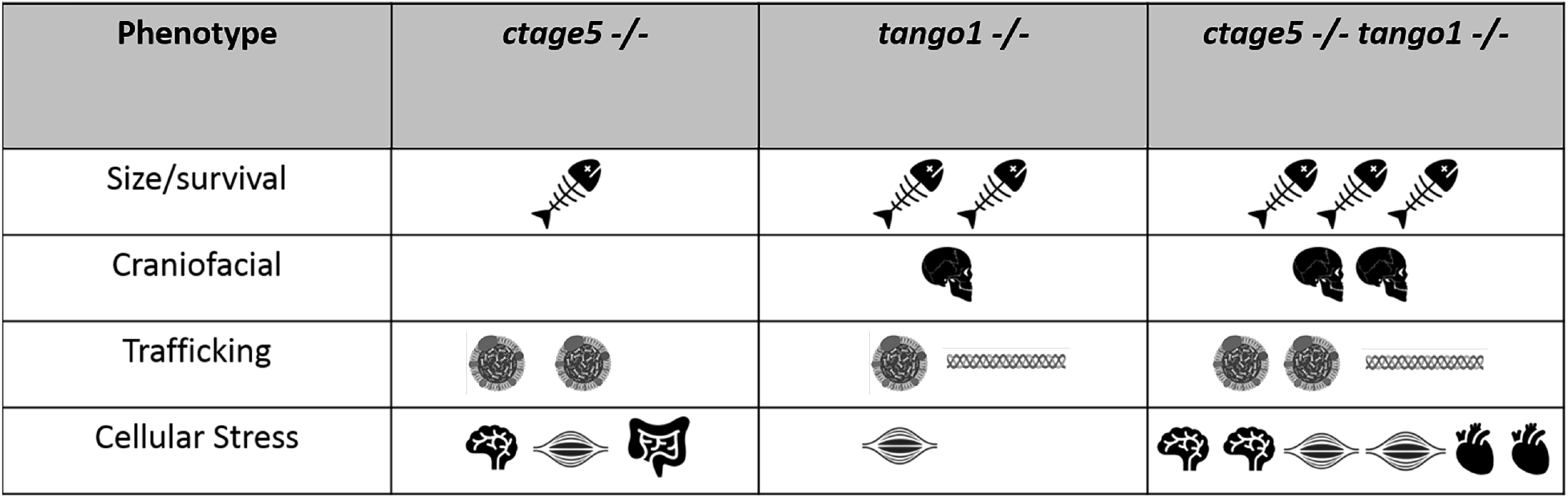
Summary of Phenotype magnitude in mutant fish. Table summarizing phenotype severity. Any symbol represents an elevated severity compared to *wildtype*. More symbols represent increased severity. 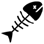 Survival/Size defects, 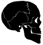 craniofacial defects, 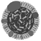 lipoprotein, 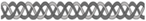 collagen, 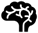 brain, 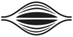 muscle, 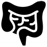 intestine, 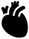 heart.

More striking differences in *tango1* versus *ctage5* mutant phenotypes were characterized for collagen secretion. Collagen II export was blocked in *tango1* mutant larvae, most notably in the chondrocytes, resulting in distended ER and craniofacial defects. *ctage5* mutants, however, have normal collagen II trafficking in chondrocytes and craniofacial structures appeared normal. These results are in line with findings from Wang et al. (2011), where trafficking of multiple collagens were affected with loss of TANGO1 resulting in severe craniofacial defects. In our study, we found similar results investigating Collagen II trafficking. Prior to our study, the role of CTAGE5 in collagen trafficking in vivo had not been investigated. In vitro studies, however, showed that collagen VII trafficking is disrupted with deletion of *CTAGE5* (Saito et al., 2011). Our results suggest that collagen II trafficking is not affected with *ctage5* deletion, indicating Tango1 has the essential role in Collagen II trafficking and craniofacial development.

In contrast to collagen trafficking, lipoprotein trafficking was predominantly affected by mutation to *ctage5*, where lipoprotein trafficking from the yolk into the zebrafish embryo was completely disrupted. Previous studies have shown that lipoprotein trafficking and lipid profiles are altered with loss of CTAGE5 or TALI (Wang et al., 2016; Santos et al. 2016, Zhang et al., 2018). Our results are consistent with these studies and extend these findings through a direct comparison of Ctage5 and Tango1 for lipid trafficking in vivo. These results suggest that Tango1 plays a primary role in collagen trafficking, whereas Ctage5 was more important for lipoprotein trafficking.

While our studies addressed the shared and unique functions between Ctage5 and Tango1 with respect to trafficking and cellular homeostasis, the experiments in zebrafish also demonstrated that Tango1 plays an important role in maintaining ER-Golgi morphology, consistent with in vitro observations (Bard et al., 2006; Rios-Barrerra et al., 2017; Reynolds et al., 2019). CTAGE5 also remodels ER-exit sites and plays a role at the ER and ER Golgi intermediate compartment (ERGIC), creating an autophagy supply pathway (Ge et al., 2017). In this regard, our observation of increased lysosomal volume in *ctage5* mutants is consistent with compensatory upregulation of lysosome-mediated protein degradation. As lysosomal volume was unchanged in *tango1* mutants, these data suggest a specific role of Ctage5 in autophagosome formation. Overall, our data confirm and extend the important, but surprisingly divergent and cell type specific roles of Ctage5 and Tango1 in large cargo trafficking and intracellular membrane dynamics.

Several observations from our studies have relevance to human disease. The nervous system was severely affected by *ctage5* deletion, consistent with descriptions in mice (Zhang et al., 2018). We found neuronal stress was not observed with deletion of *tango1* alone, but *ctage5* mutant dependent stress was exacerbated with *tango1* deletion. Interestingly, in humans, the CTAGE5 variant P521A was identified as a potential risk factor for Fahr’s disease, a rare neurologic disorder (Oliveira et al., 2007; Lemos et al., 2011). Furthermore, by investigating organ systems not been previously characterized, we discovered that cells in the intestine, muscle, and heart show elevated ER stress or morphological abnormalities in *tango1* and *ctage5* mutants. Heart morphology defects in *tango1* mutants are particularly interesting because the European Biomedical Institute GWAS catalog reports that multiple heart diseases, including coronary artery disease, coronary heart disease, and myocardial infarction are associated with *TANGO1* intron variants (Buniello et al., 2019). Very recently, homozygous hypomorphic coding mutations in human *TANGO1* was shown to result in collagen secretion defects leading to short stature, skeletal and neuronal defects, as well as insulin-dependent diabetes (Lekszas et al., 2020). Establishing the role of Ctage5 and Tango1 in multiple cellular processes and specific cell types in zebrafish, suggests this model system will be valuable to assess other candidate disease variants in human *CTAGE5* and *TANGO1*.

In conclusion, our study uses the zebrafish model system to define the in vivo consequences when the cellular functions of cTAGE5 and TANGO1 are disrupted. Through this analysis, complimentary and divergent roles for these proteins have been discovered, including identification of novel physiological processes and tissue dependences on these factors.

## Materials and Methods

### Mutant generation

Clustered regularly interspaced short palindromic repeats (CRISPR) guides were designed using ZiFiT Targeter Version 4.2 (http://zifit.partners.org/ZiFiT, **in the public domain**) against regions of zebrafish *ctage5* exon14, 5’-GGCCGCAAGGGCAGCTGATC-3’, and exon18 5’-GGCCTCCTATATGGCAAACC-3’ (ensemble transcript mia2-205). CRISPR guides targeting *tango1* were designed against exon1, 5’-TGGCTGCACCGAATAATCCC-3’ and exon7, 5’-ATCTGAAGTCGATGAGTCGG-3’ (ensemble transcript mia3-202). CRISPR gRNA templates were generated by cloning annealed oligonucleotides with appropriate overhangs into BsaI-digested pDR274 plasmid. CRISPR gRNAs were synthesized using a MEGAshortscript T7 Transcription Kit and purified using a mirVana miRNA Isolation Kit (Ambion, Austin, TX, USA). Zebrafish codon-optimized cas9 was synthesized using a mMESSAGE mMACHINE Kit (Ambion) and polyadenylated using a Poly(A) Tailing Kit (Ambion). CRISPR gRNAs and cas9 mRNA were co-injected into 1- to 4-cell zebrafish embryos from wild type ZDR fish maintained internally in the Link lab, at 10 ng/μL and 50 ng/μL, respectively, and surviving embryos were raised to adulthood before outcrossing to identify the founder fish carrying germline edits in *ctage5* or *tango1*. Offspring from these fish were raised to adulthood, then fin-clipped for genotyping (see below for details). The resulting 6285-bp deletion *ctage5* mutant and 14,043-bp deletion *tango1* mutant described here was identified via sequencing (Retrogen, San Diego, California, USA).

### Genotyping

Genomic DNA was extracted from zebrafish tissue using a Puregene Core Kit (Qiagen, Hilden, Germany). The genomic region containing either the *ctage5* or *tango1* mutation was amplified by PCR. The PCR protocol utilized primers flanking the expected deletion to identify the mutant allele and one primer flanking the expected deletion and one inside the deleted region to identify the wildtype allele.

To identify the *ctage5* wildtype allele, a forward primer for exon 13, 5’-AAGGCTCATGATAACTGG-3’, and a reverse primer for exon 14, 5’-GAGCGTTTTCTCTCTTGA-3’, creating a 171-bp amplicon. For the *ctage5* mutant allele, the exon 13 forward primer was paired with a reverse primer for exon 18, 5’-AGGAAGTGGAGGTCTAGG-3’, creating a 204-bp amplicon. To identify the *tango1* wildtype allele, a forward primer for exon 1, 5’-TGTTGGTGCTGCTATCTC-3’, and a reverse primer for exon 1, 5’-CTGCTGCATTCTTCATCCG-3’ were used to generate a 262-bp amplicon. For the *tango1* mutant allele, the exon1 forward primer was paired with a reverse primer for exon 7, 5’-GTAGTACTCACAATCTCTCC-3’, creating a 237-bp amplicon.

### Fish maintenance

Zebrafish (Danio rerio) were maintained at 28.5°C on an Aquatic Habitats recirculating filtered water system (Aquatic Habitats, Apopka, FL) in reverse-osmosis purified water supplemented with Instant Ocean salts (60mg/l) on a 14 h light: 10 h dark lighting cycle and fed a standard diet (Westerfield, 1995). All animal husbandry and experiments were approved and conducted in accordance with the guidelines set forth by the Institutional Animal Care and Use Committee of the Medical College of Wisconsin.

### Survival analysis and size measurements

Zebrafish embryos were added to the Aquatic Habitats system at 5dpf in either high density (30 embryos per tank) or low density (10 embryos per tank). At two months, zebrafish were anesthetized in Tricaine, blotted with paper towel and mass length was calculated, and fin clips were collected. gDNA was extracted from the fin clips and genotyped.

### Plastic sectioning and electron microscopy

4dpf embryos were fixed in Cacodylate fix [.1M Cacodylate, 2%PFA, 2%Glutaraldehyde (pH 7.4)] at 4°C overnight, post-fixed in 1% Osmium buffer solution, dehydrated through a series of methanol and acetonitrile washes, infiltration with Embed 812 resin (14120; Electron Microscopy Sciences, Hatfield, PA). Semi-thin transverse sections (1μm) were cut with glass knives on a Leica RM2255 microtome, heat-fixed to glass slides, stained with 1% Toluidine Blue in 1% Borax buffer, an imaged on a Nikon Eclipse E800 light microscope with an attached Sony DSC HX1 camera to capture craniofacial images. Plastic blocks were then submitted to the Electron Microscopy Facility at the Medical College of Wisconsin for electron microscopy. Ultrathin sections (70-80nm) collected using a DiATOME Ultra 45° diamond knife (MT7376; DiATOME), collected on copper hexagonal mesh coated grids (G200H-Cu; Electron Microscopy Sciences) and stained with uranyl acetate and lead citrate for contrast. Images were captured using a Hitachi H600 TEM microscope (Hitachi, Tokyo, Japan).

### TUNEL

4 dpf zebrafish embryos were fixed with 4% PFA overnight at 4°C. Fixed embryos were washed 3X PBS for 10 min, rinsed with dH20, and incubated with dH20 for 30min at room temperature followed by acetone at −20°C for 30 min and 30min dH20 to permeabilize embryos. Furthermore, embryos were washed 3X with PBST (PBS + 1% TritonX-100) and then incubated in 1mg/ml Collagenase II in PBST for 90 min at room temperature. Embryos were washed 3X for 15min in PBST, and TUNEL reaction mixture (Roche) or the label solution alone as a control was added for 2 hours at 37°C. Embryos were washed 3X 15 min in PBS, and then imaged using confocal microscopy. Mean pixel intensity was measured using ImageJ (Rasband, W.S. ImageJ, U.S. National Institutes of Health, MD), and quantified using GraphPad Prism software (GraphPad, La Jolla, CA).

### Alcian blue

Alcian Blue cartilage staining was performed similar to Hendee et al. (2018). 4 dpf zebrafish embryos were fixed in 4% PFA overnight at 4°C. Fixed embryos were washed with 1X PBS two times for 5 min, and bleached in 1ml 10% hydrogen peroxide (H_2_O_2_) in ddH2O supplemented with 50ul of 2M KOH for 1 hour at room temperature with rotation. Embryos were stained with .1% Alcian Blue solution comprised of Alcian Blue 8GX (Sigma-Aldrich, St Louis, MO) dissolved in acidic ethanol. Stained embryos were briefly rinsed with acidic ethanol and then washed 4X 60 minutes with acidic ethanol. Following washes, embryos were digested in 10μg/mL proteinase K diluted in PBST (PBS + .1% Tween20) for 1 hour at room temperature, re-fixed in 4% PFA, progressively dehydrated with ethanol, and stored in 80% glycerol. Wholemount images were obtained on a Leica MZFLIII dissecting scope with an attached Nikon E995 camera.

### Lysotracker red

7 dpf zebrafish were treated with 10uM LysoTracker™ (LysoTracker™ Red DND-99; L7528) in 1-phenyl 2-thiourea (PTU) or 1% DMSO in PTU for control embryos for 1hr at 28.5°C. Embryos were washed 3X 15 min with PTU, and then fixed with 4% PFA and imaged using confocal microscopy. Using ImageJ, a threshold was set to only identify high lysotracker staining, and the area was measured. Data was processed using Microsoft Excel (Microsoft, Redmond, WA) and graphed using GraphPad Prism (GraphPad, La Jolla, CA).

### Oil Red O

4 dpf larvae were fixed with 4% PFA overnight at 4°C. Larvae were washed 3 times with PBS, rinsed with 60% isopropanol, and then incubated in 60% isopropanol for 30 min on a rotator at room temperature. Next, larvae were dyed in fresh filtered 0.3% Oil Red O in 60% isopropanol for 3 hrs and then washed multiple times with 60% isopropanol and dH20 until the non-specific staining was removed. Washed larvae were mounted in agarose and imaged on a Leica MZFLIII dissecting scope with an attached Nikon E995 camera.

### Immunofluorescence

4 dpf larvae were fixed with 4% PFA overnight at 4°C. For notochord staining, larvae were left intact, but for chondrocyte staining trunks were removed and used for dapi staining and only the head was stained. Antigen retrieval was used for chondrocyte staining by incubating larvae in antigen retrieval buffer (150mM Tris-HCL, Ph9) followed by heating in antigen retrieval buffer to 70°C for 15min, and washed two times for 10 min in PBST (.1-1% triton in PBS) at room temperature. All following steps were completed on a rotator when possible. Larvae were permeabilized by washing once with dH_2_0 followed by dH_2_0 for 30 min at room temperature, acetone for −20°C for 30 min, dH_2_0 for 30 min, and 10ug/ml proteinase K in PBST at room temperature. 4% PFA was added for 20 min at room temperature to post-fix and washed 3X for 15min in PBST. Larvae were incubated in PBDT buffer (1% DMSO, 1% BSA, .5-1% TritonX-100, 1X PBS) with 5% Goat Serum (NGS) for 60min at room temperature. Collagen II (II-II6B3; 1:100; Developmental Studies Hybridoma Bank) was stained in the chondrocytes in PBDT/5% NGS at 4°C overnight. Larvae were washed with PBST 6 times for 15min/wash at room temperature and then incubated with secondary antibody (1:400) in 2% NGS/PBDT for 4 hours at room temperature. Larvae were washed 6 times with PBST for 15-20 min, and if necessary, were stained with DAPI (1:1000 in PBST) for 30min at room temperature and washed 3 times in PBS. Larvae were then mounted and imaged using confocal microscopy.

### Adult lens dissection

Three-month-old zebrafish were fixed with 2% PFA overnight at 4°C. Zebrafish lenses were dissected out and imaged using a Leica MZFLIII dissecting scope with an attached Nikon E995 camera.

### Labeled fatty acid injection into yolk

To analyze lipid transport from the zebrafish embryo yolk into the body, fluorescently tagged BODIPY® (BODIPY® FL C_12_; Invitrogen D3822) stock was dried down completely in a speed vacuum and re-suspended in canola oil to a final concentration of 1mg/ml. 4.6nl of the BODIPY® Oil mixture was injected into the zebrafish yolk at 4 dpf. Four hours after injection, zebrafish intersegmental vessels in the trunk were imaged using confocal microscopy. For measuring mean pixel intensity of BODIPY®- C_12_ in the intersegmental vessels, three vessels were averaged as a single data point using Microsoft Excel (Microsoft, Redmond, WA) and graphed using GraphPad Prism (GraphPad, La Jolla, CA).

### RNA extraction and real time PCR

Analysis of *ctage5 and tango1* transcripts was performed on whole embryos. mRNA was extracted from dechorionated embryos at 7dpf using Trizol-Chloroform treatment. The isolated aqueous phase from the resulting Trizol-Chloroform gradient is transferred to a new tube, and incubated with isopropanol at room temperature for 10 minutes. Centrifugation at 4°C for 10 minutes results in formation of an RNA pellet, which is then washed with 75% Ethanol in DEPC-treated water. Following this wash, the pellet is allowed to dry, and is resuspended in DEPC-treated water with a 10 minute incubation at 60°C. Resuspended RNA is then subjected to a DNAse treatment, and concentration is quantified.

cDNA was generated using the Superscript III First-Strand Synthesis System for RT-PCR Kit (Invitrogen) per manufacturer’s instructions and all qRT-PCR was performed on a CFX Connect Real-Time System (Bio-Rad) using prime time gene expression master mix (IDT). qRT-PCR was performed on five biological replicates, and all biological replicates were run in triplicate for each transcript. The zebrafish housekeeping gene *ef1a* was used for normalization. Statistical significance of differences in transcript abundance was calculated using Welch’s t-test. qPCR sequences are listed below:

*ctage5* exon 7-9:

Primer 1: CTCATCTTGGCCGCTTCTAT
Primer 2: CATCGACGGCAGCACTAATA
*ctage5* exon 19-20:

Primer 1: GGCGGAGGCATTGACATTA
Primer 2: AGAGAAGGCTCTGGAGATATGA
*tango1* exon 28-29:

Primer 1: AGAGGTCCAGGCGGAAA
Primer 2: CACACATCGGCCCGTTT

### Imaging and data analysis

Fluorescence was detected using Nikon Eclipse E600FN or C2 Nikon Eclipse 80i confocal systems. Mean pixel intensity was measured using ImageJ (Rasband, W.S. ImageJ, U.S. National Institutes of Health, MD). For measuring spinal cord cell number, spot detection software was used in Imaris (Bitplane, Zurich, Switzerland). Data was processed using Microsoft Excel (Microsoft, Redmond, WA) and graphed using GraphPad Prism (GraphPad, La Jolla, CA). An unpaired, two-tailed t test was used to analyze graphs with two groups. For three or more groups, a one-way ANOVA was conducted with Tukey’s post-hoc analysis for pair-wise comparisons.

### Adult brain clearing

PACT clearing was performed based Cronan et al., 2015 with slight modifications. Briefly, zebrafish were fixed in 2% paraformaldehyde (PFA) for 1 day at 4°C. The fixed whole adult fish brain was dissected and incubated at 4°C for 1 day in ice cold, freshly made hydrogel monomer solution of A4P0 (4% acrylamide in PBS) supplemented with 0.25% VA-044. A4P0-infused samples were incubated for 3 h at 37°C to initiate tissue-hydrogel hybridization. Hydrogel monomer solution was removed, and washed 3X with PBS. Whole brains were then incubated in 8% SDS in 200 mM boric acid, pH 8.5, at 37°C with shaking for 5 hours or until the brain periphery was clear. This solution was replaced with PBST and samples were left to shake overnight at room temp, allowing the center of the brain to clear overnight. Samples were then washed throughout the day four changes of PBS, 0.1% Triton X-100 at room temperature, and then incubated in RIMS imaging media (Yang et al., 2014) overnight at room temperature on a rotator. Samples were stored in RIMS at room temperature until imaging using confocal microscopy.

### Transgenic fish

Establishment and validation of 5XATF6RE:d2GFP transgenic zebrafish was previously described (Clark et al., 2020).

## Acknowledgements

The authors thank Michael Cliff, William Hudzinski, and Edi Kuhn for zebrafish husbandry. Jon Bostrom for help with CRISPR design, and Dr. Elena Semina and her laboratory for assistance with alcian blue staining. We are also grateful to Clive Wells for help with electron microscopy experiments.

## Author contributions

E.M.C. and B.A.L. designed experiments. E.M.C. performed and analyzed experiments. E.M.C. and B.A.L wrote the manuscript. All authors read and approved the final manuscript.

