## Supplemental Figures for "Complementary and divergent roles for Ctage5 and Tango1 in zebrafish"

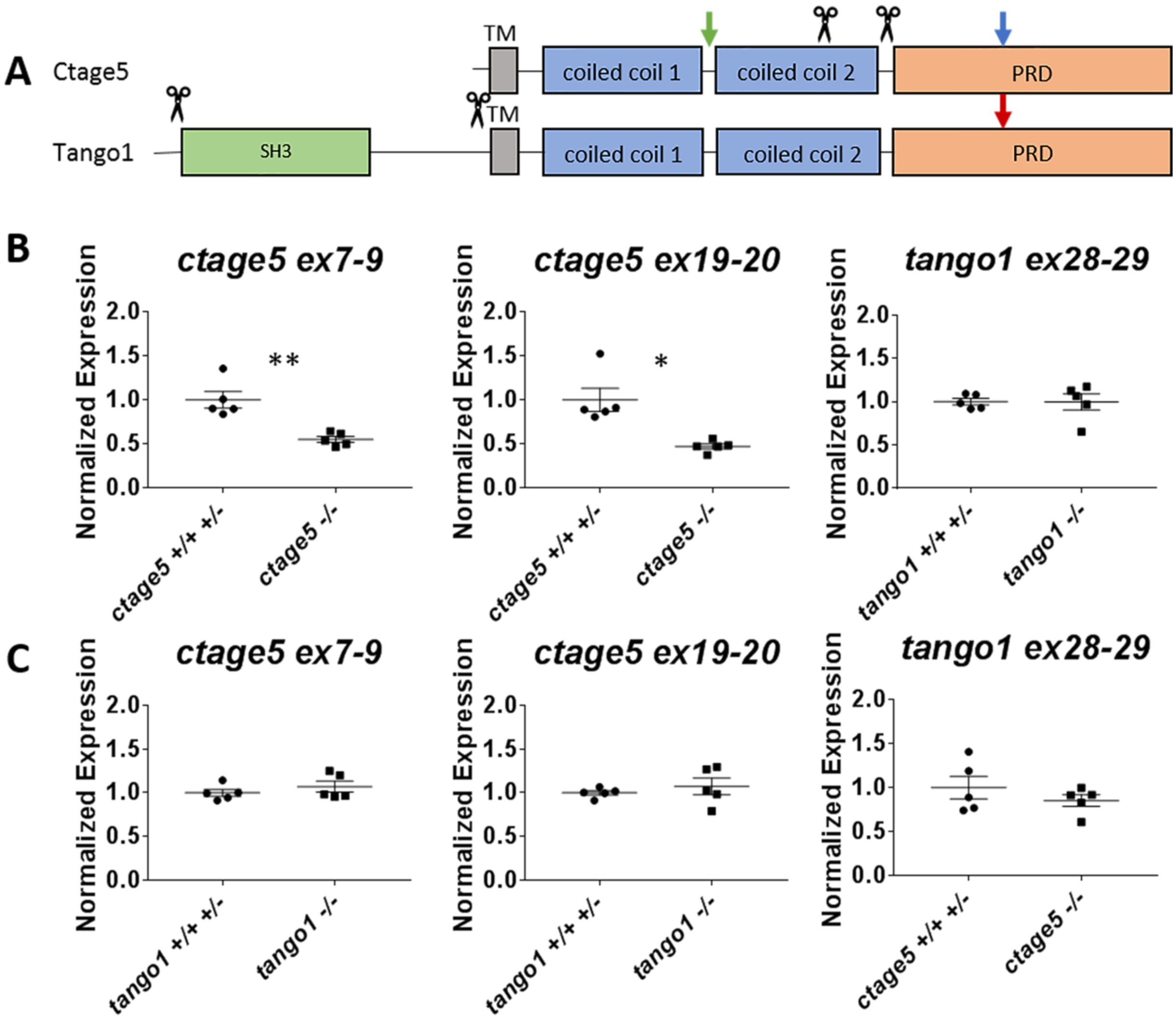


**Fig. S1: *ctage5* mutants have decreased transcript:** (A) Schematic showing large deletion cut sites (scissors), and qRTPCR primer set locations (green arrow=*ctage5* exon 7-9; blue arrow=*ctage5* exon 19-20; red arrow=*tango1* exon 28-29). (B) qRTPCR t-test results investigating alterations in gene expression in 7 dpf large deletion mutants using the specified primers. Welch's t-test was used to analyze statistical differences (*ctage5* ex7-9, p=0.0061; ctage5 ex19-20, p=0.0147; *tango1* ex28-29, p=0.9716) (C) qRTPCR results investigating compensation of *ctage5* or *tango1* gene expression in 7dpf *tango1* or *ctage5* large deletion mutants respectively using the specified primers. Welch's t-test was used to analyze statistical differences (*ctage5* ex7-9, p=0.3762; *ctage5* ex19-20, p=0.4954; tango1 ex28-29, p=0.3534). *=p<.05**=p<0.01.

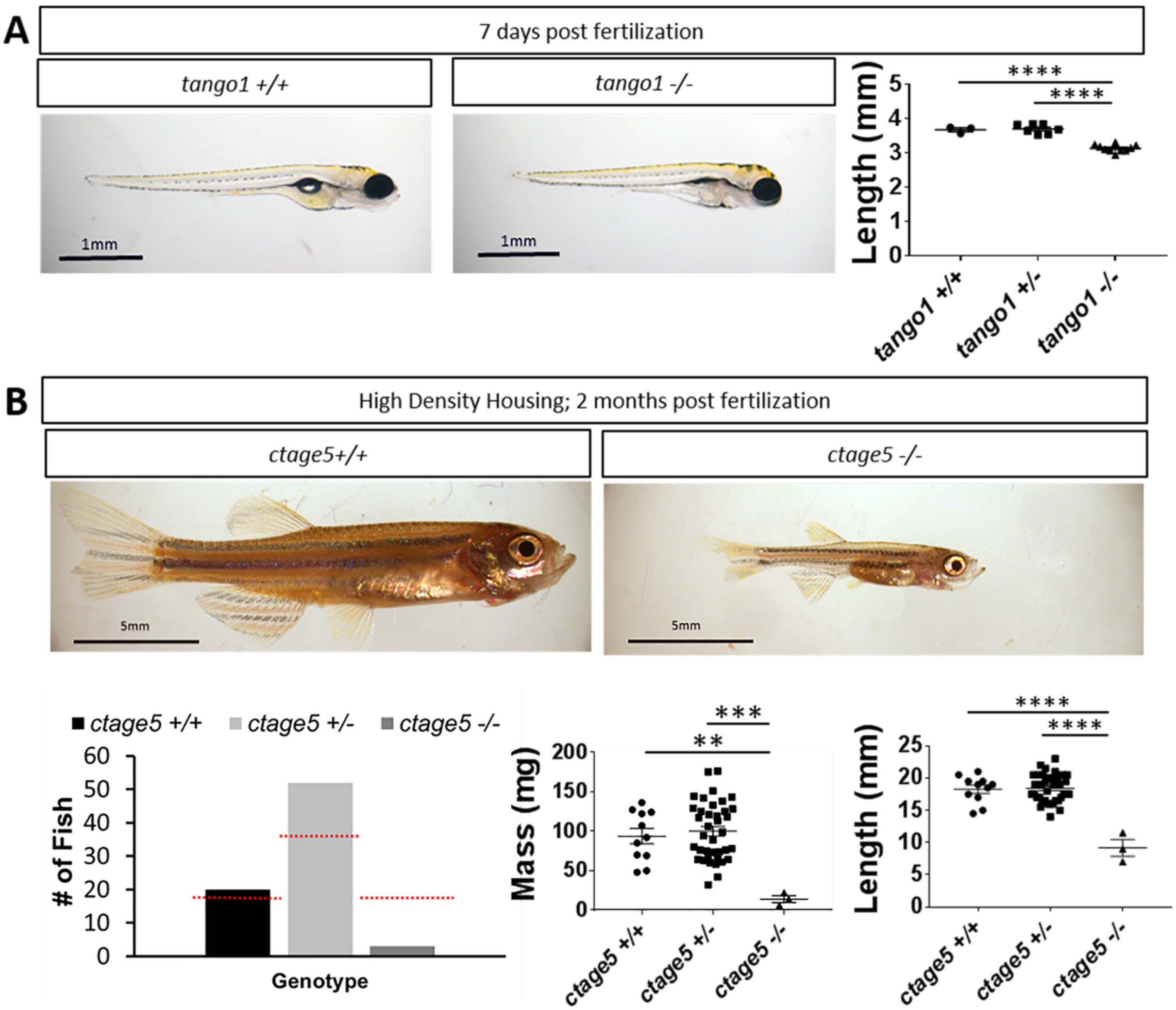


**Fig S2: Size differences in *ctage5* and *tango1* mutants.** (A) Length measurements for 7dpf embryos (one-way ANOVA, F=55.95, p<0.0001). (B) Representative images of *ctage5* +/+ and *ctage5* -/- 2-month-old zebrafish raised in a high density environment (about 30 fish per tank) and quantification for survival ($\chi2$=18.92, p<0.05, n=75), mass (one-way ANOVA, F=8.372, p=0.0007), and length (one-way ANOVA, F=26.69, p<0.0001). Red-dotted lines in represent expected survival, and bars are actual survival.
